# Calculating sample size requirements for temporal dynamics in single cell proteomics

**DOI:** 10.1101/2020.12.09.418228

**Authors:** Hannah Boekweg, Amanda J. Guise, Edward D. Plowey, Ryan T. Kelly, Samuel H. Payne

## Abstract

Single cell measurements are uniquely capable of characterizing cell-to-cell heterogeneity, and have been used to explore the large diversity of cell types and physiological functions present in tissues and other complex cell assemblies. An intriguing application of single cell proteomics is the characterization of proteome dynamics during biological transitions, like cellular differentiation or disease progression. Time course experiments, which regularly take measurements during state transitions, rely on the ability to detect dynamic trajectories in a data series. However, in a single cell proteomics experiment, cell-to-cell heterogeneity complicates the confident identification of proteome dynamics as measurement variability may be higher than expected. Therefore, a critical question for these experiments is how many data points need to be acquired during the time course to enable robust statistical analysis. We present here an analysis of the most important variables that affect statistical confidence in the detection of proteome dynamics: fold-change, measurement variability, and the number of cells measured during the time course. Importantly, we show that datasets with less than 16 measurements across the time domain suffer from low accuracy and also have a high false-positive rate. We also demonstrate how to balance competing demands in experimental design to achieve a desired result.

## Introduction

Individual cells express a unique proteome; this is true for cells in a complex environment as well as cells in a laboratory-controlled cell culture experiment. These differences arise from both intrinsic and extrinsic factors, such as access to nutrients, spatial relationships to other cells or cell cycle status^1^. Tissues in multicellular organisms often contain a variety of discrete cell types, each of which expresses a unique proteome, and the combination of these functional cell states gives rise to the overall tissue function. Single cell measurements facilitate the study of such differences by quantitatively measuring transcript or protein abundances for individual cells^2,3^.

Much of the early work in single cell phenotypic characterization was done by mRNA sequencing^4,5^, and this still remains a widely used data source. However, intra- and extracellular functions are most often carried out by proteins, e.g. cytoskeletal structures, metabolic enzymes, signal transducers, etc. Previous work over the past decade has revealed that mRNA abundance measurements are a poor proxy for protein abundance measurements, reviewed in^6–8^. Indeed, many important temporal trends in the dynamic proteome are not detected in mRNA data^9^. This is true both for bulk measurements and also for single cells^10–12^. A recent investigation of proteome dynamics during the cell cycle found only 15% of mitotic cycling proteins had coordinated cycling mRNA transcripts^1^. Thus, to identify dynamic proteome responses at the single cell level, proteomic measurements are necessary.

Single cell proteomics poses an enormous technical challenge and, until recently, global proteome profiling had not been demonstrated, reviewed in^13^. As single cell proteomics gains momentum, it is important to note the practical limitations that are encountered for its practitioners, particularly as they relate to the number of cells that can be analyzed within an experiment. Single cell mRNA-sequencing experiments, which benefit from ligation-based barcoding^14^, are able to multiplex tens of thousands of samples into a single data acquisition run^15^. However, proteomics multiplexing remains limited to ~20 samples^16^. Therefore, the primary limitation in the proteomic profiling of a very large number of individual cells (>1000) is still instrument acquisition time. For this reason, many researchers face practical limitations in the type of experiment that can be designed to investigate proteomic phenotypes at the single cell level.

Characterizing proteome dynamics over time is critical for understanding cellular differentiation, disease progression, and treatment response. In contrast to a two-state comparison, a time-course experiment collects measurements several times during a biological process. One fundamental question for this experimental design is how many time points need to be sampled in order to detect protein dynamics. Because of the practical limitations of single cell proteomics, it is possible that the number of time points analyzed may not be sufficient to achieve statistical confidence in the dynamic trends. Whereas the T-test commonly used in a two-state comparison is well-behaved with a small number of samples in each state (e.g. 5-10), it is unknown how such a small sampling will impact the ability to detect trends in time-course data. A variety of tools have been created to help map expression dynamics, or trajectories^17–19^. However, these are often created within the assumptions of an experiment where the number of sampled cells across the time domain is potentially thousands. Herein, we present a method to facilitate estimating the number of cells needed in a temporal dynamics experiment by systematically exploring the impact of proteome variability and effect size through a large simulation similar to a power analysis.

## Methods

All calculations and data used in this manuscript can be found in our publicly available GitHub repository, https://github.com/hboekweg/SingleCellSampleSize. Below are listed specific scripts used to create figures and metrics cited in the manuscript.

### Calculating Accuracy and False Discovery

For the data presented in Figures 1 and 2, we simulated a large population of ‘cells’ with a single protein abundance and time measurement using the standard formula y=mx + b as protein_abundance = slope * time + 1 + ε. The error term, ε, is a random error which mimics the biological and/or technical variability in a measurement. This error is drawn from a zero-centered Normal distribution with a specified standard deviation. For a single simulation, slope and standard deviation are chosen from: slope s ∈ [0.5, 1, 2, 4], standard deviation v ∈ [0, 0.25, 0.5, 0.75, 1]. We then create a population with random seeds for the time variable between 0 and 1. The full software for making simulated populations can be found in the GitHub repository in a file called simulate_data.R.

**Figure 1 -.**
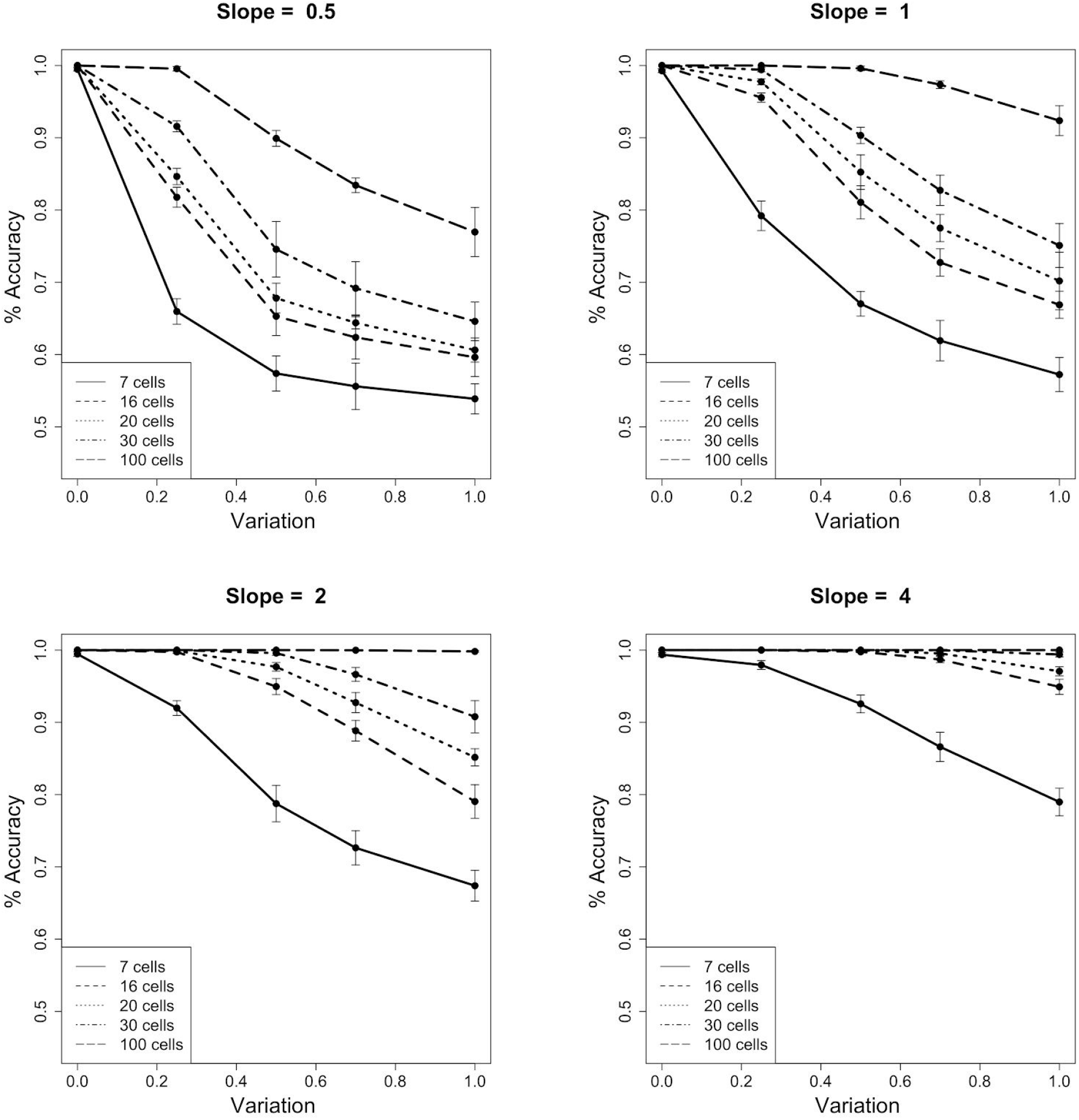
Accuracy in the identification of temporal dynamics. For various parameter sets of slope, variability and number of cells, the accuracy of correctly identifying temporal dynamics is shown. For comparison, each of the four subpanels is organized by slope, and shows a parameter sweep over equivalent measurement variabilities and cell numbers. Error bars are derived from 10 independent simulations.

**Figure 2 -.**
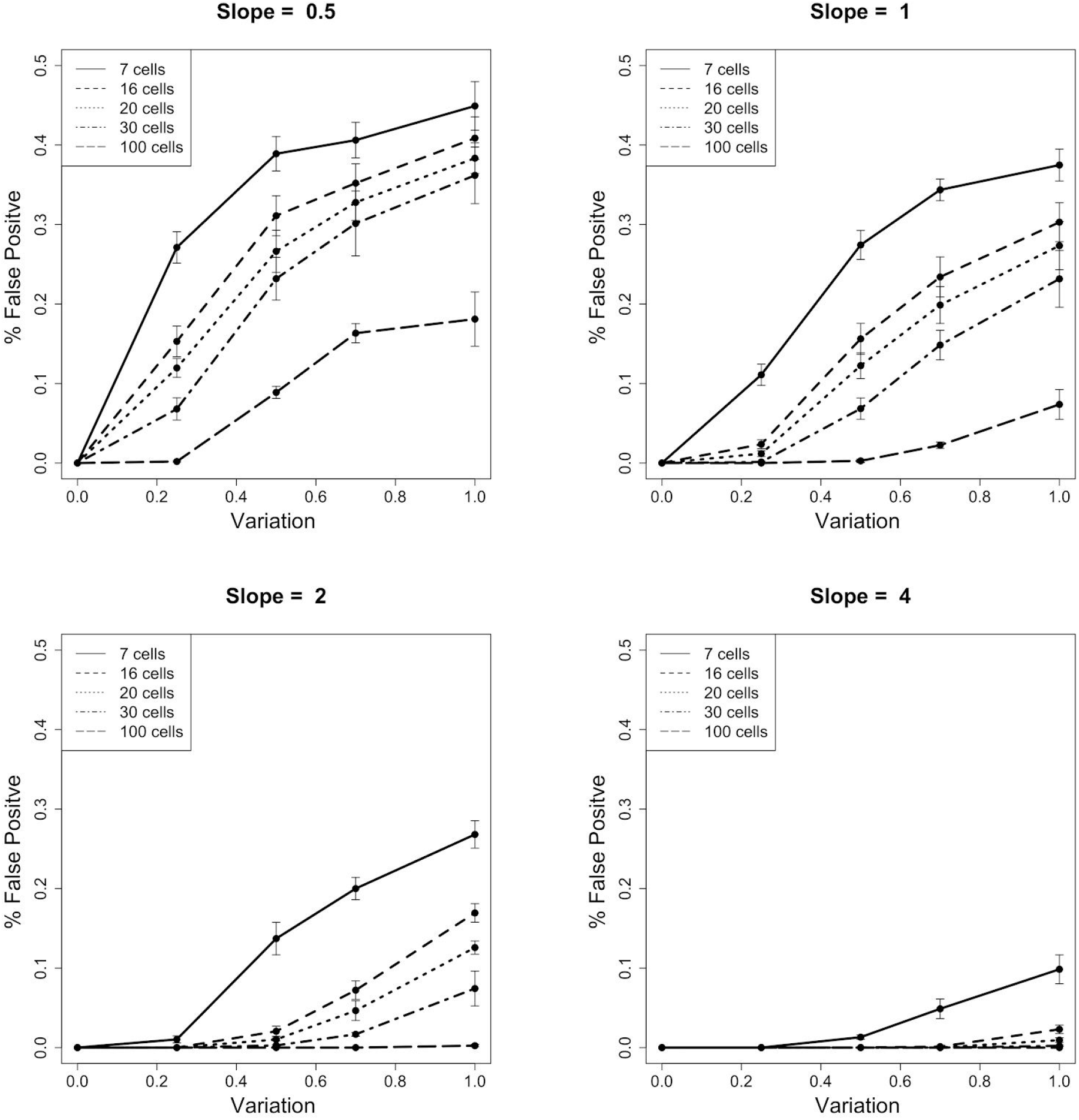
False-positive rate in the identification of temporal dynamics. The rate of false-positive identification was calculated for the same set of parameters seen in Figure 1. False-positive is defined as the misclassification of a non-changing protein, falsely reporting it as changing. As seen in Figure 1, panels are organized by slope, and show the parameter sweep across variation and number of cells.

To determine the true-positive and false-positive rates, we sampled a specific number of cells from the population, n_sample ∈ [7, 16, 20, 30, 100]. With the protein abundance and time values for these subsampled cells, we used cellAlign^17^ to interpolate a trajectory (Supplemental Figure 1). Next, we calculated the area between the interpolated trajectory and the population’s true temporal trajectory, ABC_true. The null hypothesis (no change) was tested by calculating the area between the interpolated trajectory and a horizontal line at the mean protein abundance of the subsampled cells; this metric is called ABC_null. If the ABC_true < ABC_null, then we assert that the data from the subsampling represents a changing protein; if ABC_true > ABC_null, then we assert that the subsampling represents a non-changing protein. The accuracy reported in Figure 1 is the result of 1000 sub-samples per population. We also repeated the entire simulation 10 times, which is shown on Figure 1 as the error bars. The full software for calculating accuracy and making the panels of Figure 1 can be found in our GitHub repository in the file called makeFigure1.R.

The process for calculating the false discovery was very similar to calculating accuracy, except that the correct answer was that there was no change, and the incorrect answer was that interpolated trajectories matched a sloped line. We simulated a large population of cells, using the same formula as above, with the caveat that this population always had a true slope of zero. To calculate false discovery, we again subsampled cells from the population and interpolated a trajectory using cellAlign. We then calculated an area between the curve metric ABC_true which reflected the distance between the interpolated trajectory and the population’s true trajectory (i.e. no change). We also calculated an ABC_falsepositive metric, which reflected the area between the interpolated trajectory and a sloped line. For the plots in Figure 2, the slope was either 0.5, 1,2, or 4. The false positive rate reported in Figure 2 is the result of 1000 samples per population. We repeated the entire simulation 10 times, which is shown on Figure 2 as the error bars. The full software for simulation, calculations and plotting the panels of FIgure 2 can be found in our GitHub repository in the file makeFigure2.R.

### Generalizing Slope and Variation

To generalize the true-positive and false-positive rates, we ran a separate simulation to re-calculated these metrics using the slope/variation ratio and number of subsampled cells. The data for Figure 3 was generated using the same types of simulations as described above for Figures 1 and 2 except that different slopes and variations were used to calculate S/V values of 0.5, 1, 1.5, 2, 3, 4, and 6. The plotted data comes from different simulations of a single S/V ratio with different values for S and V respectively. For example, data for S/V of 0.5 was generated using S/V = (0.5/1.0; 0.75/1.5; 1.0/2.0; 1.5/3.0; 2.0/4.0; 3.0/6.0). The other S/V values were calculated in the same manner. The full code used to simulate, calculate and plot data for Figure 3 can be found in our GitHub repository in the file makeFigure3A.R and makeFigure3B.R.

Supplemental Figure 2 re-analyzes published single cell proteomics data^20^, and uses two cell types, C10 and SVEC, to plot fold change between cell type and the within-group variation. The fold change for each protein is found by taking the absolute difference between the mean abundance for C10 cells and SVEC cells. The plotted variation is the standard deviation of C10 cells. The full code used to analyze, calculate and plot data for Supplemental Figure 2 can be found in our GitHub repository in the file makeSupplementalFigure2.R.

**Figure 3 -.**
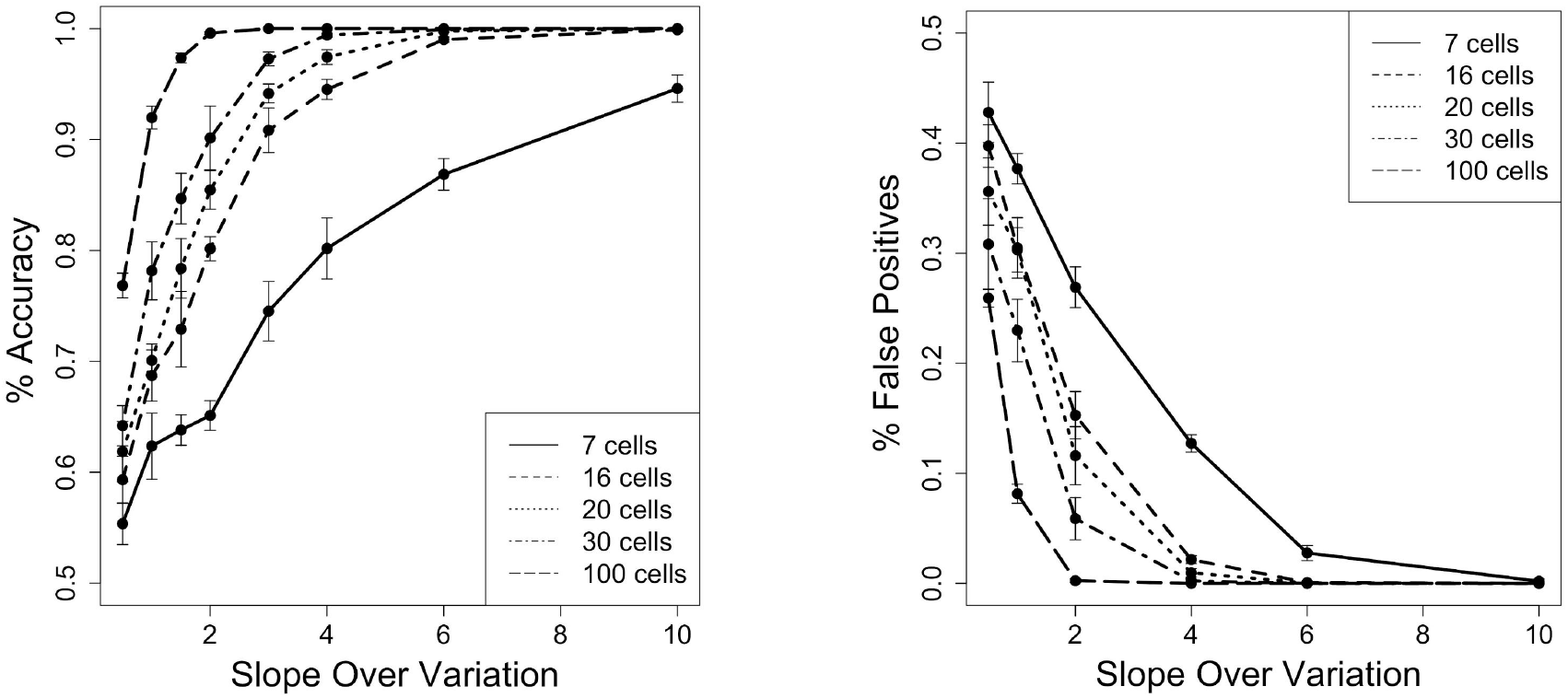
Scale invariant trends. Accuracy and false-positive rates are shown for a scale-free ratio of slope/variation. Similar to Figures 1 and 2, simulations are used to determine the true-positive and false-positive rates of various parameter combinations of slope, variation, and number of cells. A specific slope/variation datapoint is derived from multiple different combinations of slope and variation. For example, values plotted for S/V=0.5 were derived from six simulations using slope/variation = (0.5/1.0; 0.75/1.5; 1.0/2.0; 1.5/3.0; 2.0/4.0; 3.0/6.0). Note the y-axis scale is zoomed to allow better visualization of the data.

### Estimating S/V

To characterize how well a subsample of measurements can approximate the true slope and variation of a larger population, we ran a simulation as described above with true S/V = 1. From the large population of 5000 cells, we subsampled a given number of cells n_sample ∈ [7, 16, 20, 30, 100], and calculated S/V_est_. The S/V_est_ is calculated by using linear regression to fit a line to the subsampled data. The slope of the fitted line is then used as our estimated slope. The estimated variation is found by calculating the standard deviation of the residuals off the fitted line. We calculated S/V_est_ for 1000 independent subsamples of the population and computed the distribution of the difference S/V_true_ - S/V_est_. This histogram is plotted in Figure 4A. Figure 4B shows the number of proteins that would remain after using S/V_est_=1 as a cutoff. We simulated 5 populations with 1000 cells, at S/V_true_ ∈ [0, 0.5, 1, 1.5, 2]. We subsampled 30 cells from a population, calculated S/V_est_ and discarded the subsampling event if S/V_est_ < 1. The figure shows the percent of sampling events that remained after filtering. The full code used to analyze, calculate and plot data for Figure 4 A and B can be found in our GitHub repository in the file makeFigure4.R and simulate_data_figure4.R.

**Figure 4 -.**
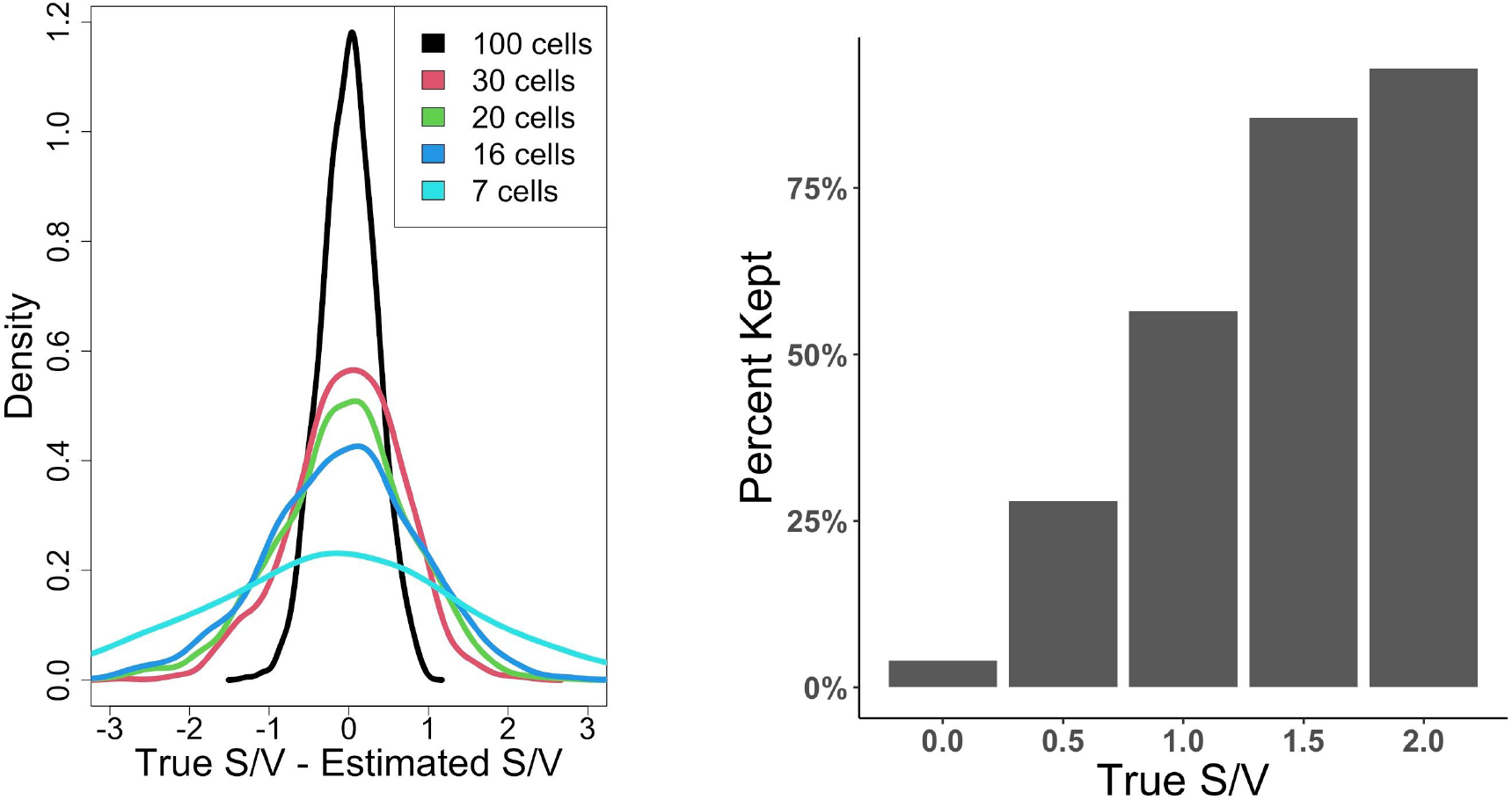
Estimating accuracy of S/V approximation of data. (A) We estimated the S/V from a subsampling of cells, where the true population S/V = 1. The histogram shows the difference between the approximated S/V and the true S/V, using subsample sizes of 7, 16, 20, 30, and 100. (B) The effect of using an estimated S/V as a cutoff. Data was simulated to contain proteins with S/V of 0, 0.5, 1, 1.5, and 2. After removing data with S/V_est_ < 1, the graph shows the percentage of proteins kept according to their true S/V.

## Results

One of the challenges associated with mass spectrometry measurements is that they are inherently destructive. In order to measure the proteins in a cell, the cell itself is destroyed (e.g. lysed). Thus, for an experiment that measures change over time, the cells measured at time_0_ will be different from the cells measured at time_1_. This means that the change in protein abundance observed between time_0_ and time_1_ has at least two potential sources. Some of the change can be attributed to temporal dynamics that are a deterministic part of the experiment brought about by the perturbation. The second source of change between cells measured at time_0_ and time_1_ is attributed to cell-to-cell variability. We specifically emphasize that even for synchronized cells, or experimental designs where great effort has been taken to homogenize cellular state, there will be a real and observable variability in the proteome between individual single cells. Furthermore, this variability is often much greater than one might initially expect.

A second important element of a time-course experiment is how time is measured. There are numerous relevant applications of time in biology, including: cellular differentiation, the transition from health to disease, or response to an external stimulus. In these various experimental systems, time can be absolute (2pm) or relative (5 minutes after stimulus). Time can be an observable fact (date) or a value inferred from the data (relative time during the cell cycle as measured by the abundance of various markers). In studying disease progression, the time metric is more accurately a ‘pseudotime’ that measures the approximate progression from a healthy to a diseased state - perhaps measured by visible morphological features. Depending on the specific scientific question, an experimental sampling of the time domain might be rigorously defined at very specific intervals or a random sampling. In this manuscript and the statistical simulations described below, time is abstract. The variable representing time varies within the bounds from zero to one; zero represents the temporal beginning of an experiment and one represents the end.

### Simulations

To understand how measurement variability, effect size and the sample size affect our ability to detect temporal changes in proteome dynamics, we performed a large simulation across the relevant parameter space. Each simulated cell has one protein measurement and an associated time variable, calculated from two input parameters. First, the simulation uses a simple linear change as the shape of temporal dynamics, and in our model this rate of change is called the slope. Second, we specify a measurement variability modeled as a normally distributed error term. Using slope, variability and a time value between 0-1, we can calculate a protein abundance (see Methods). The simulation starts by creating a large number of cells which cover the entire time range; each cell is represented as an [abundance, time] data point. From the large simulated population, we subsampled a small number of cells with the primary goal of determining whether this small sample accurately represents the larger population. The subsampled measurements are used to interpolate a temporal expression trajectory using cellAlign^17^; this interpolated trajectory was then compared to the true population trajectory and a null model (see Figure S1). If the subsampled trajectory was closer to the true population trajectory, then we classified this subsampling event as correct; if the subsampled trajectory was a better fit to the null model, then this subsampling event was classified as incorrect.

We ran multiple simulations with different parameter values for slope and variability. In each simulation we calculated the average accuracy (see Methods). Figure 1 shows the accuracy of detecting protein dynamics for various combinations of slope, measurement variation and the number of sampled cells. Several expected trends emerge from the simulation. First, the accuracy improves if more cells are sampled across the time domain. Regardless of the slope or measurement variability, increasing the number of cells improves the temporal expression trajectory interpolated from the subsampling. The smallest number that we sampled, 7 cells, has poor accuracy in almost any [slope, variation] parameter set. A significant improvement is seen in increasing from 7 to 16 cells, followed by steady improvement as the number of cells increases to 20, 30 or beyond. Second, accuracy decreases with greater measurement variability. For example, in the simulations where slope = 1, a 16 cell sample has an average accuracy of 95% if the measurement variability is 0.25, but has an average accuracy of 68% if the variability = 1.0.

To characterize the potential for incorrectly labeling a protein as changing, when it actually was unchanged in the time course, we simulated data where the slope parameter was zero. From this large population, we again subsampled, interpolated an expression trajectory using cellAlign and compared the trajectory to the population’s true trajectory (i.e. no change) or a simple sloped line. If the interpolated trajectory more closely matched a sloped line, despite being composed of data points in a population whose slope was zero, we counted this as a false-positive. As we discovered in simulations above, the number of sampled cells and the measurement variability have a significant impact on the false-positive rate (Figure 2). In several simulations with a small subsampling (e.g. 7 cells), the false-positive rate approached 40%.

### General Principles

To help generalize the results of our simulation, and make them more immediately applicable to proteomics datasets, we grouped parameter sets by their slope/variation ratio. The simulations in Figure 1 and 2 report results which used a convenient numeric scale. However, proteomic datasets reported in the literature have a wide range of possible values, with some datasets reporting raw quantitative values in the millions and others using log-transformed, zero centered data. By transforming our results into a slope/variation ratio (S/V), we directly test whether the observed true-positive and false-positive trends are scale-free. Thus, regardless of how quantitative protein data is obtained or processed, the S/V ratio can provide applicable guidance.

For a wide range of slope/variation ratios, we ran simulations to generate the true-positive and false-positive rates (see Methods). This explicit exploration of the relationship between expression change and variability revealed distinct trends in the ability to correctly identify protein dynamics. For example, in an experiment with 16 cells, an S/V = 2 would have an 80% true positive rate (Figure 3A), meaning that if 100 proteins had this trajectory slope and measurement variability, we expect to detect 80 of them (the other 20 go undetected). Under the same conditions, we also see a 15% false positive rate. This means that of all the non changing proteins, 15% of them would be incorrectly identified as changing. As with the results presented in Figures 1 and 2, the true-positive and false-positive rates improve when a larger number of cells are sampled across the time-course. If an experiment sampled 30 cells instead of 16 cells, the true-positive rate would improve from 80% to 90% for S/V =2; coordinately the false-positive rate would fall from 15% to 5%. These charts are essential in understanding how the accuracy of temporal trajectory detection depends both on the number of cells analyzed across the time domain and also the trajectory’s slope and the protein’s inherent measurement variability.

To help frame these results, we sought to understand S/V values for proteins in real data. We examined a single cell proteomics experiment which compared two different cell types^20^, with a sufficient number of replicates to obtain a reliable estimate of within-group variability (n>20). We note that this dataset does not demonstrate temporal dynamics, but rather the magnitude of differences in protein abundance between biological states. This can still be used to approximate slope if time is scaled to a unit value. Fold-change and within-group variability were calculated for each protein in this dataset (Supplemental Figure 2). Most proteins have a small fold-change, and often the magnitude of fold change is similar to the magnitude of variability. Thus, relatively few proteins have a S/V ratio above 1. As shown in Figure 3, proteins with an S/V below 1 will have a low true-positive and high false-positive rate unless a very large number of cells are used. Proteins with a more attractive true-positive and false-positive rate, such as S/V = 2, are only ~3% of proteins in the dataset. Proteins with S/V > 4 are exceedingly rare (< 0.2%).

### Without an Oracle

The analysis of real data does not benefit from knowing the true protein expression dynamics, as one knows in a simulated dataset. Thus, when trying to apply the lessons learned in a simulation (e.g. Figure 3) to a real world time-course dataset, researchers must estimate slope and variation of a protein in their data. We investigated how well we can estimate the slope and variation (S/V_est_), using the same simulation methodology as above. We calculated the precision of S/V_est_ for sample sizes of 7,16, 20, 30, and 100 cells (Figure 4A). As expected, the precision of this estimate improves when more cells are samples. When we sample 100 cells, the S/V_est_ is typically very close to the real value; the standard deviation of the error is 0.34. If we sample 30 cells, the standard deviation of the error is 0.67. To demonstrate the effect of using this estimate, we simulated how many proteins of various S/V values would be removed from a dataset if S/V_est_ was used as a filtering criteria (Figure 4B). If an experiment sampled 30 cells across a time-course and used a S/V_est_ = 1 for a cutoff, approximately 92% of high quality proteins (S/V = 2) would be retained.

### How Many Cells?

A proposed set of dynamic proteins discovered in a time course experiment represents a mix of true-positive and false-positive identifications. Although the statistical simulations above may help estimate the relative rates of false-positives, it is not possible to point out which specific identifications are potentially suspect. The best way to clarify this list and winnow out false-positives is via replicate analyses. If the same experiment is independently replicated, the expected true-positive rate can be expressed as *p^x^*, where *p* represents the probability and *x* represents the number of independent replicates. For example, in a 30 cell time-course experiment with two replicates, we would expect the true-positive identification rate for proteins with S/V = 2 to be 0.9^2^, or 0.81; the expected false-positive rate would be 0.05^2^, or 0.0025. Predicting the true-positive or false-positive rates for a more complex experimental design, such as requiring *n* observations among *k* replicates, can be determined using standard statistical sampling methods. Using these expected rates and the relative number of proteins for various S/V values suggested from Supplemental Figure 2, scientists can appropriately plan for various experimental design scenarios.

A challenging part of experimental design is balancing competing priorities and appropriately budgeting a limited resource. As previously discussed, the primary limiting factor in a single cell proteomics study is the total number of cells that can be analyzed. This number is currently much less than desired, which forces researchers to choose whether more cells should be devoted to a single time-course or to additional replicate time-courses. As a simple illustrative example, imagine allocating a budget of 50 cells in a time course experiment (Figure 5). Option A involves two replicates with 25 cells each; option B allocates 16 cells into each of three replicates. With option A, researchers achieve a more dense sampling of the time-course and therefore have better true-positive and false-positive rates per replicate. Option B has an additional replicate. Although the rates for true-positive and false-positive are not as good in each individual time course in option B, leveraging three replicates drives the overall false-positive rate lower than in option A.

**Figure 5 -.**
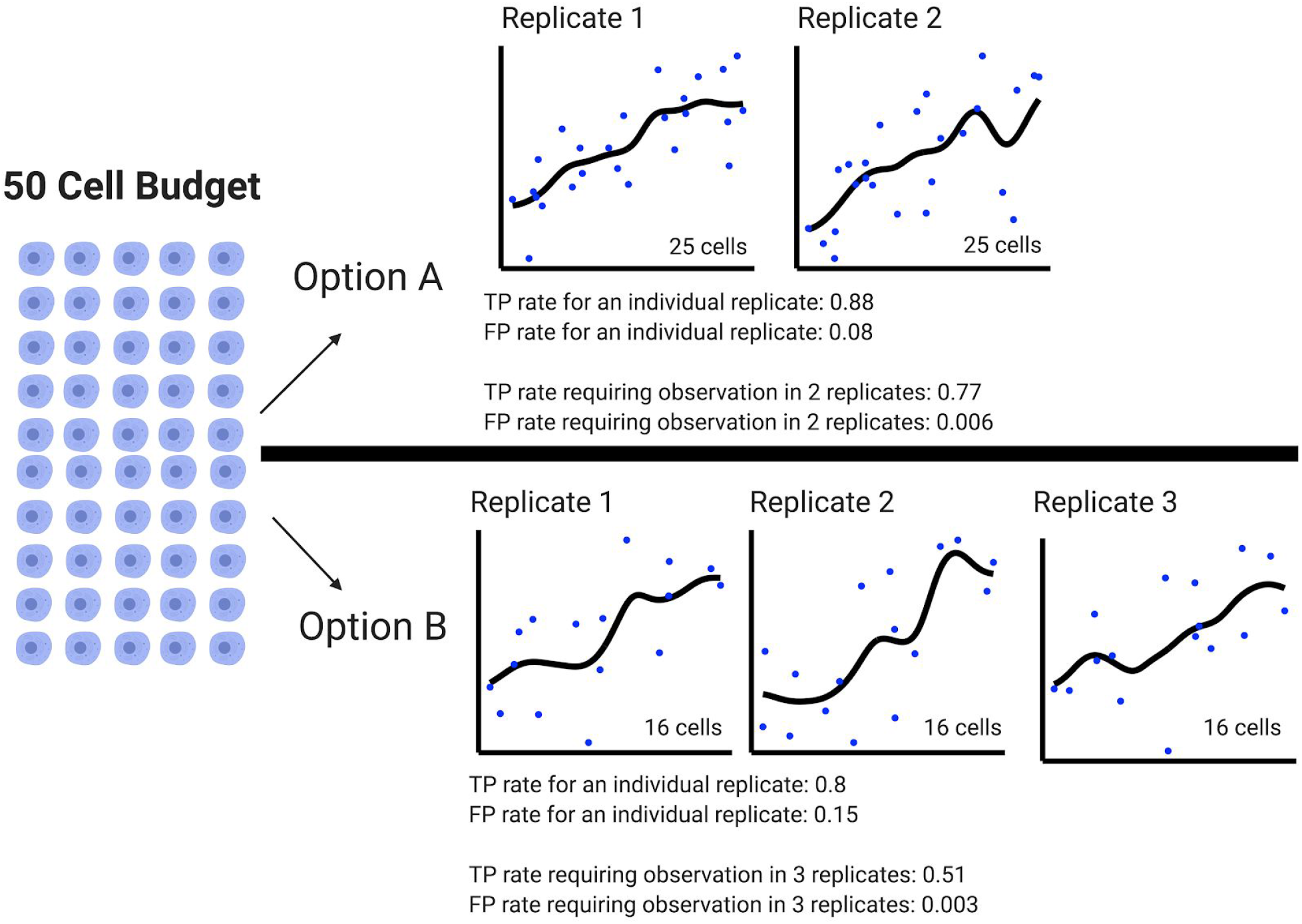
Scenarios for allocating a limited number of cells. With a total budget of 50 cells, two different options are demonstrated. True-positive and false-positive rate for each individual time course and the overall rate with replicates is shown. Option A depicts an experiment with two replicates and 25 cells characterized during each time course. Option B shows an experiment with three replicates and 16 cells characterized during each time course. When considering replicates, option A has a higher TP rate but option B has fewer false positives.

## Discussion

Single cell proteomics is an emerging technology that promises to help clarify the diversity of cellular phenotypes as well as reveal essential trends in proteome dynamics. The current throughput of single cell proteomics is significantly lower than single cell sequencing technologies. Therefore, until there is a dramatic change in proteomics instrumentation and associated technologies, the proteomics community will struggle with the necessity of analyzing fewer cells than they would like. Within this context, it is essential to properly plan experiments on a limited number of cells to maximize the likelihood of success. Statistical power calculations exist for two-state comparison experiments that rely on the t-test. However, temporal trajectory experiments lack an experimental planning tool to help estimate the accuracy of different designs. Here we have simulated the accuracy of detecting a temporal change in protein abundance against the null hypothesis of no-change. Simulations explored a variety of metrics such as the magnitude of temporal change (slope), cell-to-cell heterogeneity (variability), and the number of cells analyzed across the time domain. The simulations highlight the need to analyze a sufficient number of cells across a time-course; a higher number of cells across a time-course always leads to more favorable true-positive and false-positive rates.

All projects work within the bounds of a budget, and herein we discussed a budget as the number of cells that can be analyzed. Choosing a specific experimental design is a balancing act and may require compromises between the desire to have enough cells to accurately approximate the temporal expression trajectory and practical limitations of the budget. In the broad array of experiments conducted in biomedical and environmental sciences, investigators will have to make this difficult choice. In a clinical experiment monitoring drug response, patient demographics may be particularly compelling, and demand more patients from diverse backgrounds. With a fixed budget of cells, the choice to analyze more patients reduces the number of cells analyzed for each patient. Alternatively, the biological sample itself may be a limiting factor. For rare cell types or highly degraded/diseased samples, there may be a finite number of cells that are available to be analyzed - regardless of budget. Finally, we note that the experimental process itself might make it difficult to obtain a desired sampling density within the time domain.

Our statistical simulations were motivated by their utility in the emerging field of single cell proteomics. However, the results are applicable to any analysis attempting to identify dynamic trajectories, such as longitudinal studies of an individual person over time^21,22^. For these and similar studies, it is important to understand the variability of a measurement. We show that most proteins have a within-group variability equivalent to or greater than the observed fold change between conditions. If variability to this degree is present in the data, it will be very challenging to confidently detect temporal dynamics with a limited number of measurements across time. Even for studies which utilize rigorous clinical assays with defined technical variability, biological variability must be anticipated and characterized.

Finally, we note that the simulations herein model only simple linear increases and not more complex expression patterns. A common biological experiment measures the response to an external stimulus, often reporting a temporary change followed by a return to the original state^23^. A classic example of this is transient phosphorylation signaling. Yet other biological investigations monitor cyclical expression changes related to light/dark patterns and circadian rhythms^9,24^. Based on our results, we expect that detecting these complex non-linear patterns will require a dense sampling of the time domain.

## Acknowledgements

This work was funded through a sponsored research agreement from Biogen Inc. The authors declare no financial conflict of interest.

---

**Figure S1.**
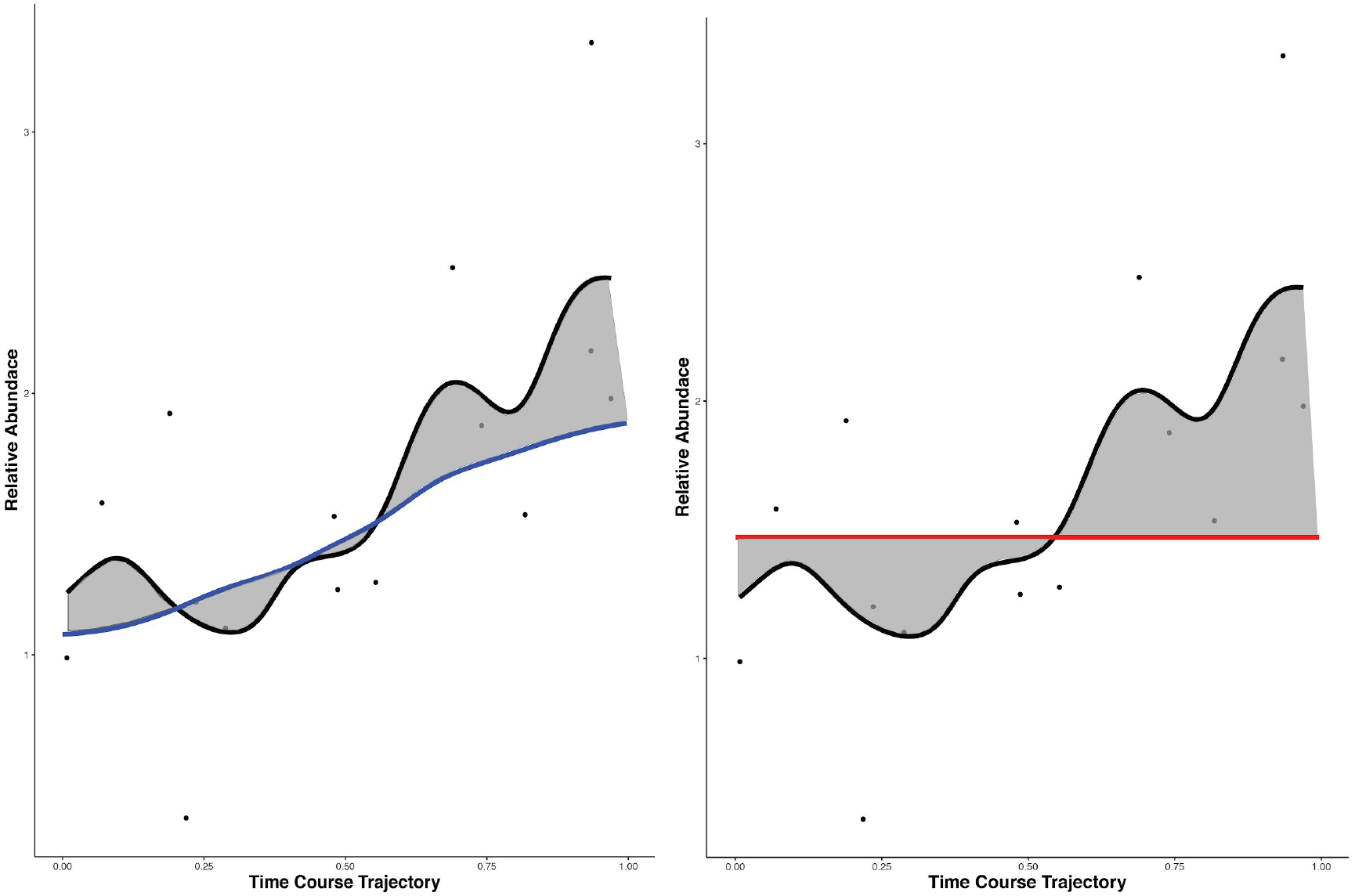
Subsample interpolation and ABC calculation. Both panels show a 16 cell subsampling from a larger population of measurements made with slope = 1 and standard deviation = 0.5. The trajectory of these 16 data points is interpolated with cellAlign and shown in a black line. In the left panel, the subsample’s interpolated trajectory is compared with the true trajectory derived from the original population (blue line). The area between these two lines is calculated as the ABC_true (shaded in grey). In the right panel, a null model of no-change is evaluated. The subsample’s trajectory is compared to a flat line equivalent to the average intensity value; this metric is called ABC_null.

**Figure S2.**
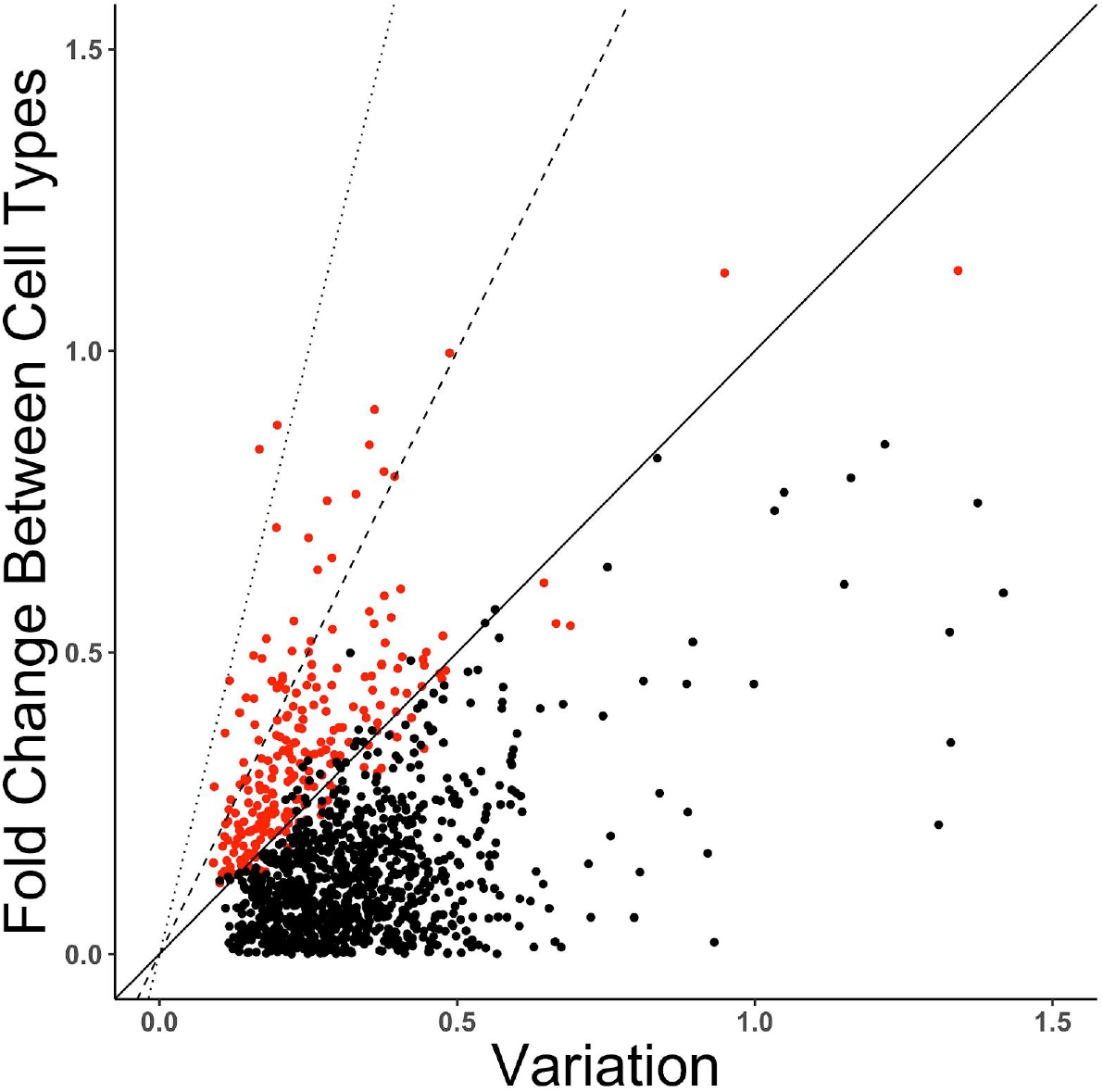
Fold change versus variability of proteins in a single cell proteomics dataset. Using quantitative proteome abundance data^20^, we calculated the standard deviation of within-group replicates and compared this to the fold change between cell types. Each dot on the graph is a different protein, and red indicated that the protein would have passed a T-test for differential expression (FDR corrected p<.05); proteins not passing a T-test are shown in black. For convenience, we have drawn lines indicating a 1:1 ratio between fold change and variation (solid line), a 2:1 ratio (dashed line) and a 4:1 ratio (dotted line).

